# Influence of selected plant growth stimulators in enhancing germinability and germination parameters of *Zea mays* L. under microgravity conditions simulated by a two-dimensional clinostat

**DOI:** 10.1101/2021.11.23.469749

**Authors:** Beckley Ikhajiagbe, Geoffrey O. Anoliefo, Alexander O. Orukpe, Saheed I. Musa

**Affiliations:** Environmental Biotechnology and Sustainability Research Group, Dept. of Plant Biology and Biotechnology, University of Benin, Benin City, Nigeria; Applied Environmental Biosciences and Public Health Research Group, Department of Microbiology, University of Benin, Benin City, Nigeria; Space-Earth Environment Research Laboratory, University of Benin, Edo State, Nigeria; National Space Research and Development Agency, Obasanjo Space Complex, Abuja, Nigeria; Centre for Atmospheric Research, Anyigba, Kogi State University, Nigeria; Department of Biology and Forensic Science, Admiralty University of Nigeria, Delta State, Nigeria

**Keywords:** Clinostat, microgravity, *Zea mays*, indole acetic acid, gibberellic acid, ascorbic acid

## Abstract

The earth has become increasingly overcrowded as a result of rapid urbanization and population growth, predicting that its carrying capacity could be overstretched. As a result, it is important to test the possibilities of growing plants under space exploration conditions, especially gravitational balance. Since microgravity impedes plant development, to what extent can plant growth stimulators reverse or enhance this trend? A total of 12 maize seeds were weighed and placed sideways in petri dish and inoculated with plant growth stimulators such as indole acetic acid (IAA), gibberellic acid (GA), and ascorbate (AA) and the clinorotated at different rates (0.5, 1.0 and 2.0 rpm), while control seeds were just placed on a table. Results showed that at 72 hrs, the maize seeds under microgravity showed reduced germination percentage with increasing clinorotation rates as against the control. But when stimulated with IAA, GA and AA, improved germination percentage was observed as against the control even under microgravity condition. The seedling dry weight, germination time and other germination parameters also showed similar improvements. Comparatively, the three growth stimulators showed no major variations in their ability to improve germination percentage under microgravitational impact. However, IAA showed more improvement on seedling vigor as against others, while GA showed more effect on the peak time and rate of germination. This research confirmed the possibilities of improving germinability of maize seeds under space exploration condition.

## 1. Introduction

As the planet becomes increasingly overcrowded as a result of accelerated urbanization and population growth in all countries, regardless of their level of technological or economic development, there has recently been a movement to find alternative places for humans to live (Vandenbrink et al., 2014). The concern is that the earth’s carrying capacity would be exceeded, and the planet will be unable to meet the world’s ever-increasing population’s needs for food, nutrients, oxygen, and other resources. Scientists have started looking for alternative habitation sites in order to avoid a potential humanitarian catastrophe. This has aided research into alternative planets to see what possibilities there are for human habitation in the event of a humanitarian crisis (Braun et al., 2018). However, it’s also worth noting in the field of space science that space exploration can be difficult to accomplish before resources are available in sufficient quantities (Raymond, 2017).

According to Soga et al. (2002), Humans on board the spacecraft will need metabolic energy in addition to the energy required to power the spacecraft. Traveling through space could take years, and since humans are heterotrophs, resources such as oxygen for metabolic energy production and energy in the form of food would be needed. By the way, these two forms of human sustenance criteria for space travel are all plant-based. Plants, for example, have long been thought to be the best oxygen generators, considering the use of synthetic methods (Stutte et al., 2002). Plants use up CO_2_, which is a by-product of human metabolism, and then give off oxygen, which is needed for human survival. Photosynthesis is also very important, especially in driving the process for the generation of ATP, which humans require for metabolism (Papaseit et al., 2000). Second, the primary producers in the food chain, where humans are at the top, are plants (Kering and Zheng, 2015), so plants cannot be taken away from humans’ survival, whether as food, during the production of oxygen for ATP formation, or as an electron carrier during cellular energetics.

The Earth’s gravity, which exerts a force of “1 g,” has resulted in the formation of plants. During space travel, one of the greatest effects on plants is the influence of gravity (Levine, 2010; Vandenbrink et al., 2014; Braun et al., 2018; Kiss et al., 2019; Orukpe et al., 2021). This ubiquitous force influences plant development, productivity, and morphology at all levels, from the molecular to the whole plant (Vandenbrink et al., 2014). Studies have shown that changes in gravitational effects usually affect development of plants. Other physical phenomena governed by gravity include buoyancy, convection, and sedimentation, all of which have an effect on a number of physical and chemical processes, as well as plant growth and development. Buoyancy, for example, affects gas exchange, cellular respiration, and photosynthesis, but it is also a characteristic of densities (Braun et al., 2018).

Studies have also shown that prolonged exposure to microgravitational effects by astronauts usually impair their metabolism (Morrow et al, 2005). The fact that this impair human metabolism is also a pointer to the fact that perhaps it may also impair plant metabolism (Ikhajiagbe and Musa, 2020). Changes in gravity have been shown to affect protein metabolism and, of course, protein synthesis (Koryum and Chapman, 2017). Since proteins are needed for the regulation of cell activities and the control of metabolic processes, anything that affects protein production, hinders or improves such processes, can eventually affect metabolic processes or metabolic process regulation. In the absence of buoyancy-dependent convective transport, reduced mass transport and thicker boundary layers around plant organs can also affect these processes in microgravity (Monje et al., 2000).

Indole Acetic Acid (IAA) and Gibberellic Acid (GA) are plant growth stimulators that can manipulate a range of growth and developmental phenomena in a variety of crops. Plant height, number of leaves per plant, and fruit size have all been found to increase with IAA, resulting in increased seed yield (Kapgate et al., 1989; Lee, 1990). Gibberellic Acid (GA) influences many aspects of plant growth and development, including growth, flowering, and ion transport, while Ascorbic Acid (AA) plays a regulatory role in promoting productivity in many plants (Shah, 2004). Ascorbic acid controls phytohormone-mediated signaling, as well as a variety of physiological processes in plants and stress protection, by acting as a cofactor for a variety of enzymes (Barth and Mario, 2006, Smirnorf and Wheeler, 2000, Farooq et al., 2013). These growth stimulators have also been recorded to help plants survive in stressful situations. The big question is whether soaking maize seeds in these chemicals before exposing them to simulated microgravity would improve their chances of survival. Soga (2002) reported the possibility for Growth stimulation of plants under microgravity conditions in space.

In a previous report, microgravitational effect on maize seeds under 0.5, 1.0, and 2.0 rpm clinorotation significantly decreased maize seed growth capacities and germinability (Orukpe at al., 2021). The maize germinants’ enzyme status was also significantly impacted. The current research aims to see if the stress effects on maize germinants can be alleviated by using growth stimulants like IAA and GA, as well as an antioxidant like AA.

## 2. Materials and methods

### 2.1 Location

The experiment was conducted at the Space-Earth Environment Research Laboratory, University of Benin operated by Centre for Atmospheric Research of the National Space Research and Development Agency (NASRDA).

### 2.2 Maize seeds

The maize seeds were obtained from the New Benin Market, Benin City, Edo State, Nigeria.

### 2.3 Preparation of Agar

Preparation followed standard methods as used by Orukpe et al. (2021). The Agar was prepared by weighing 1.5g in 100 ml of distilled water. The solution was then heated for 10mins with continuous stirring to achieve a homogenous solution. After a clear solution was observed, it was allowed to cool for another 10mins after pouring it on the petri dishes label g and µg, sowing followed immediately.

### 2.4 Sowing

The maize seeds were weighed singly using a digital weighing balance, Model No. NBT-A200 and inserted into the petri dishes containing the cooled agar, with the seeds lying sideways. A total of 12 seeds were contained on each petri dishes, and then sealed properly. The petri dish containing the control was left under the influence of gravity on a table, while the seeds meant for microgravity stimulation were placed on the clinostat for 72 hrs under different rotations (0.5 rpm, 1.0 rpm and 2.0 rpm).

### 2.5 Exposure of seeds to growth stimulators

The maize seeds that were primed in the selected growth stimulators before they were eventually exposed to clinorotation for 72 hrs. The seeds were pre-treated with 150 ppm Gibberellic acid, 150ppm Indole acetic acid and 150ppm Ascorbic acid respectively. Seeds were soaked in the above treatment for 90 mins before sowing following Musa and Ikhajiagbe, 2021.

### 2.6 Clinorotation

The experiment was prepared in two batches. The first batch were placed in clinoratation at 0.5, 1.0 and 2.0 rpm and studied for 72 hours, while the second batch were first exposed to growth stimulation in indole acetic acid, gibberellic acid and ascorbate respectively, before subjecting them to clinorotation at 0.5, 1.0 and 2.0 rpm in the 2-D Clinostat (Dimension clinostat, UN, New York, USA) (Plate 1). In the both batches, the control seeds were placed on the laboratory bench for 72 hours.

### 2.7 Germination parameters

Plant germination parameters measured were time to first germination, root length, shoot length, number of prominent roots, wet weights of seedling, dry weight of seedling and those of seedlings were measured using the weighing balance after the day of termination of the experiment. Their respective dry weights were also measured after sun drying 72 hrs. All seed germination parameters were calculated following (AOSA, 1983)

**Plate 1:**
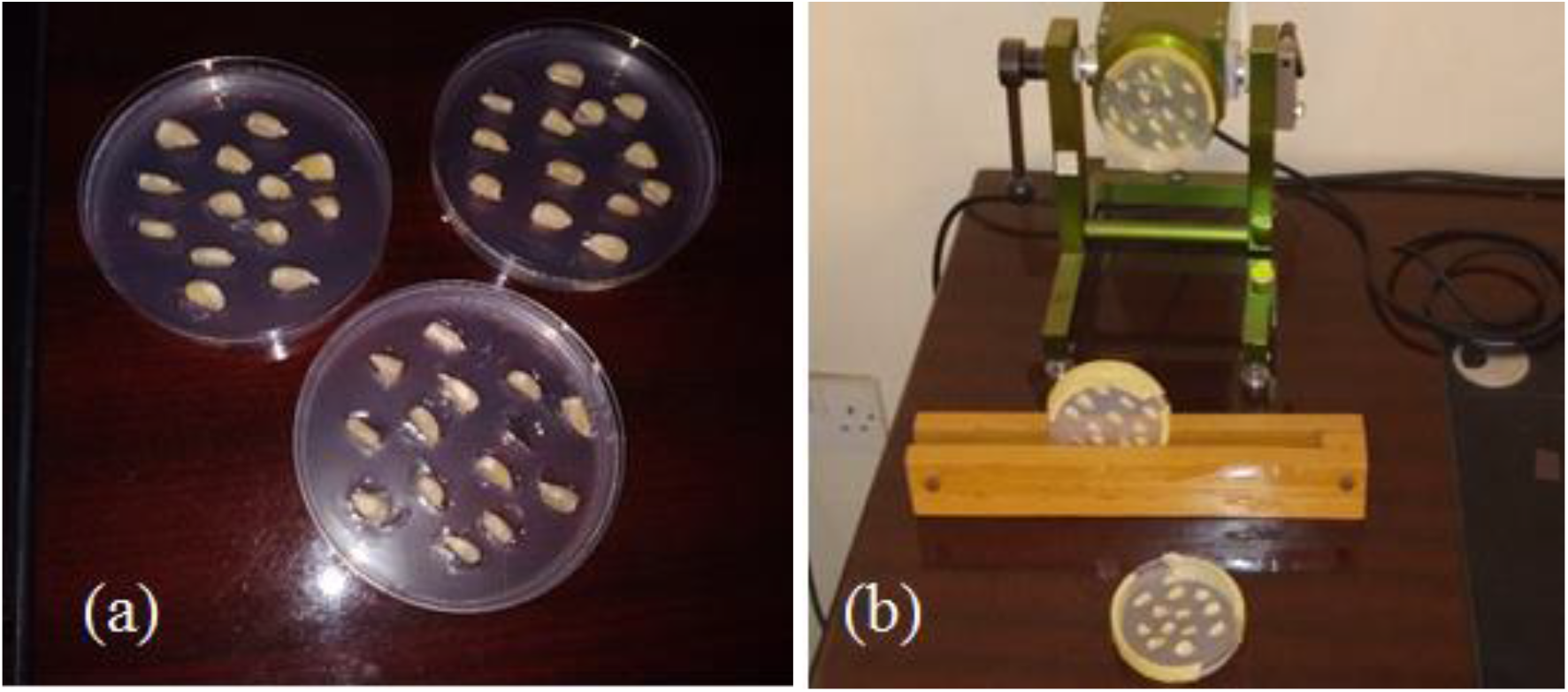
(a)The arrangement of seeds in petridish before (b) placement unto the clinostat

### 2.8 Statistical Analysis

Assuming homogeneity of the experimental set-up, a complete randomized project design was adopted. Means of 3 replicates were presented and means separated by aid of the Duncan Multiple Range Test. Germination indices were computed according to the methods laid down by Edwards (1932), Al-Mudaris (1998), Farooq et al. (2005), Ranal and Santana (2006), ISTA (2015), Aravind et al. (2020), including final germination percent, median germination time, and seedling vigor.

## 3. Results

### 3.1 Effect of microgravity on germination parameters before plant stimulation

The effect of gravity on the final germination percent of *Zea mays* and seedling dry weight (g) before exposure of plant growth stimulators is depicted in Figure 1. The final germination percent at 72 hrs was 60 % at 2.0 rpm in the clinostat, 68 % at 1.0 rpm and 80 % at 0.5 rpm compared to 94 % in the control. A similar results was observed in the seedling dry weight. This results showed a significant decrease in germination percentage and seedling dry weight as the clinostat rate increases.

**Figure 1:**
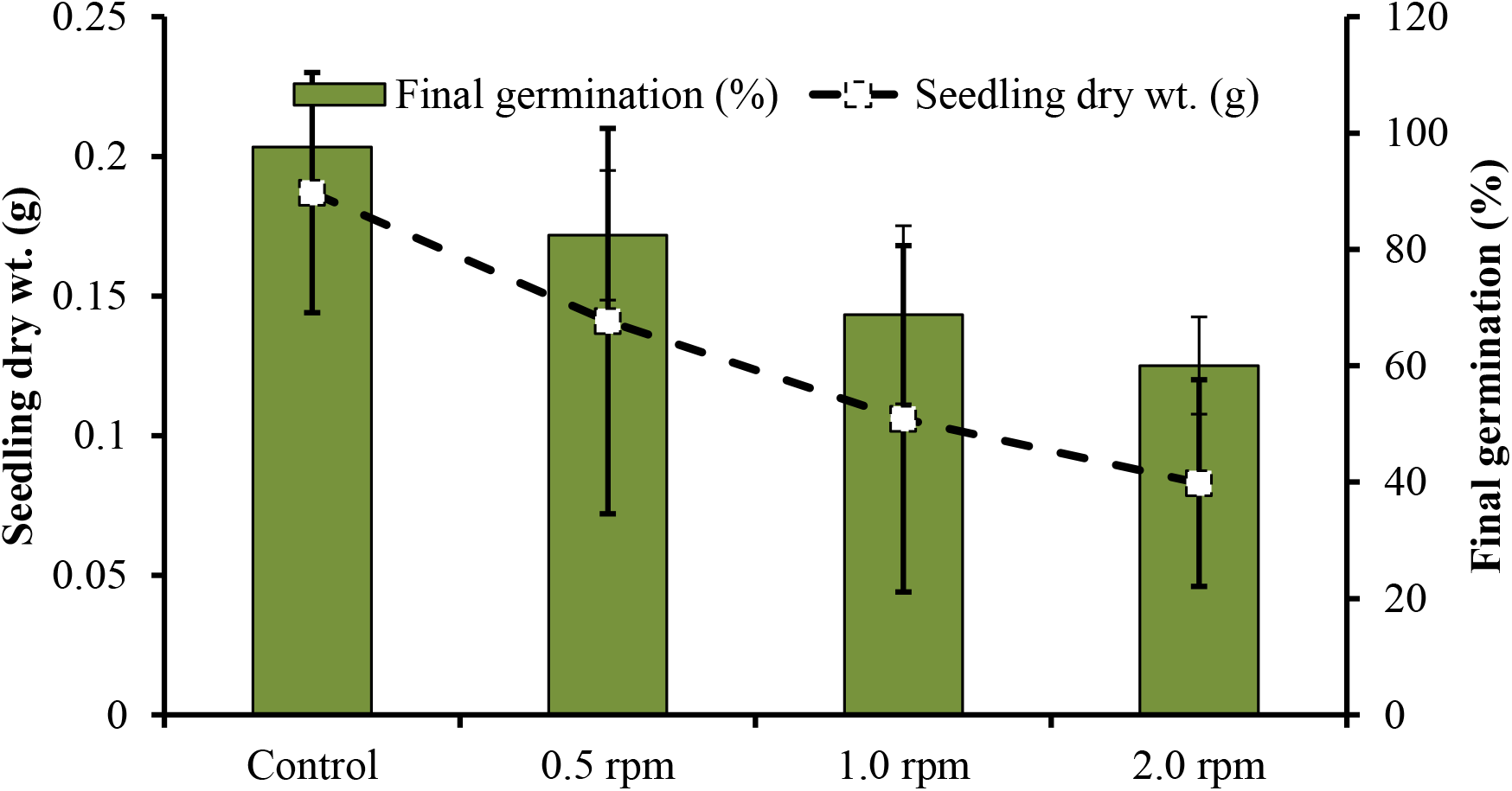
Effects of microgravity on the final germination percent and seedling dry weight of *Zea mays* at 72 hrs after germination initiation.

The time it took for seeds exposed to microgravity to germinate for the first time was recorded (Figure 2). The result showed that when seeds were exposed to microgravity, the germination significantly delayed in all the clinostat rates (0.5 rpm = 45 hr, 1.0 rpm = 54 hr and 2.0 rpm = 55 hr). However, in the control petri dish where seeds were not subjected to microgravitational influence, it took only 40 hrs for seeds to germinate. This showed that germination was delayed by 15 hrs under the microgravitational impact at 2.0 rpm.

**Figure 2:**
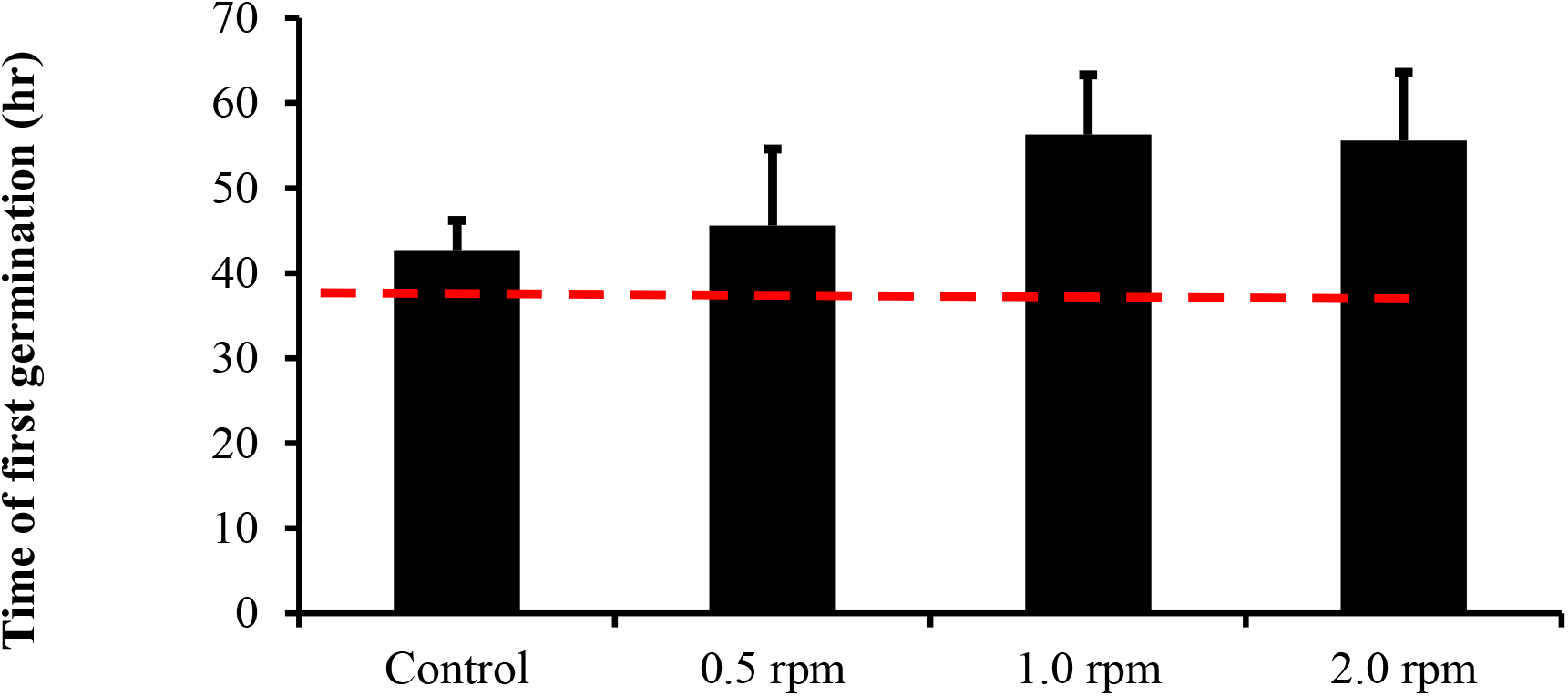
Time taken to attain first seed germination in seeds following exposure of maize seeds to micro gravitational effects

### 3.2 Effect of microgravity on germination parameters after exposure to plants growth stimulants

Gibberellic acid (GA) was added to the maize seeds before exposure to microgravity to test its effectiveness in improving maize seed germinability and germination parameters (Table 1). At first, the GA improved germination parameters at gravity compared to the control (H_2_O). The GA induced microgravity-exposed seeds germination percent at 72 hrs showed significant variations from the control (92.98-97.45 %, p<0.05). Even under microgravity, the GA-induced seeds germinated earlier than the control (H_2_O). However, no significant change in germination time was observed with the microgravity exposure at different rates. Median germination time (T50) was observed to be shorter (24.05 hrs) at the GA induced seed under gravity while an increase in T50 was observed with higher rate of clinostat. Daily germination speed was improved with the introduction of GA with no significant reduction with microgravity-induction. Generally, results showed higher effects with GA induction and slow reductions with increasing clinostat levels.

**Table 1:**
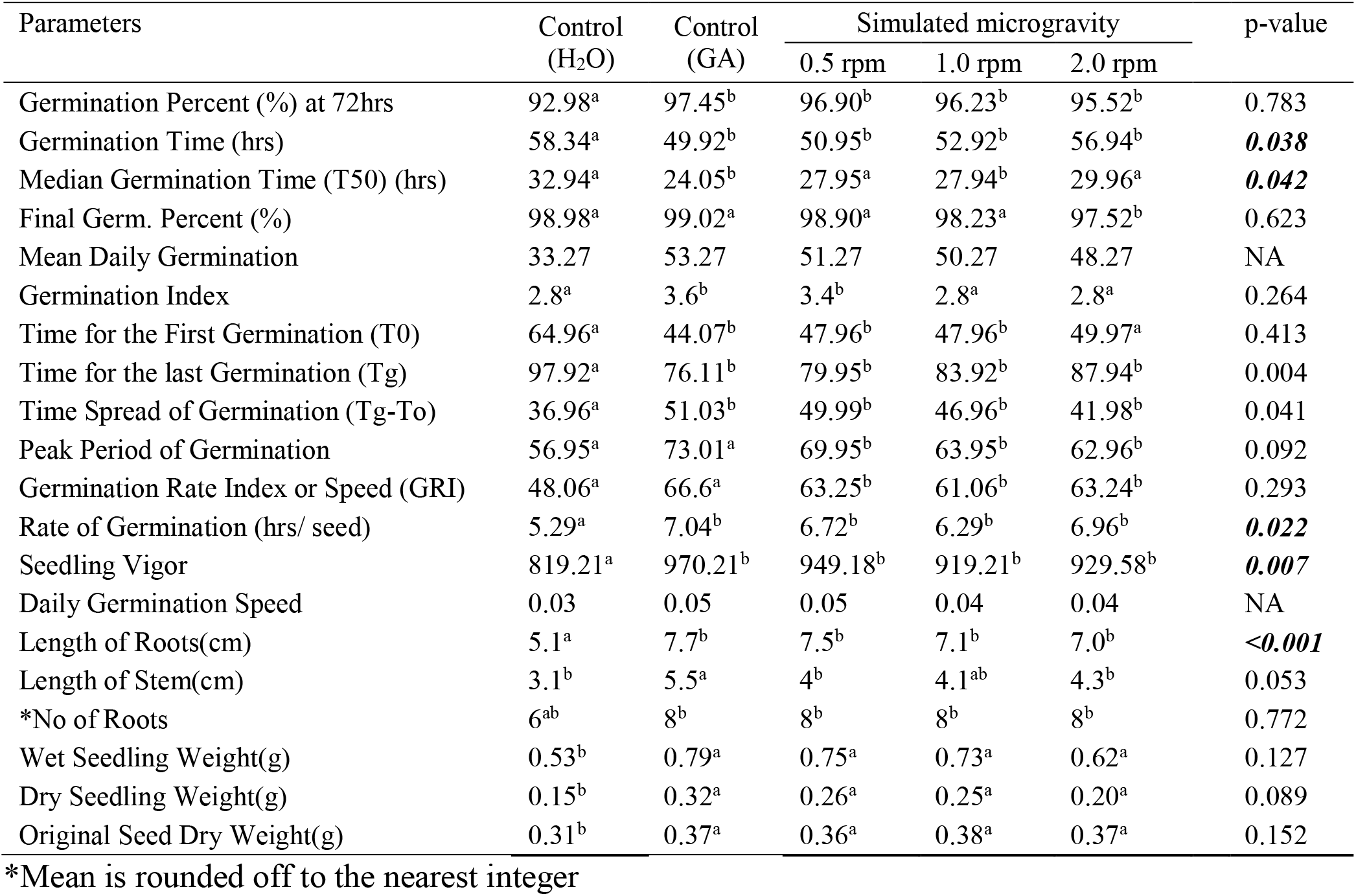
Effects gibberellic acid on the germinability and germination parameters of *Zea mays* after exposure to microgravity

When maize seeds were stimulated with IAA (Table 2), improved germination percentage was observed even under microgravity condition. The first germination appeared at 35.97 hrs in the IAA stimulated seed under 0.5 rpm clinostat rate, while the least was observed in the Control (H_2_O) (Plate 2). The peak germination period was 45.96 - 53.95 hrs (p> 0.05), according to Table 2. The germination rate index, also known as germination speed, ranged from 41.06 to 66.54 hrs (p>0.05). Microgravity-exposed seeds previously exposed to indole acetic acid had a significant increase in wet seedling weight (0.69 - 0.81g), compared to 0.5g of wet seedling weight when seeds were not exposed to microgravitational effects (control). However, when the seeds were exposed to indole acetic acid (0.15g-0.41g), there were no major variations in dry weight in relation to the degree of simulated microgravity (p>0.05).

**Table 2:**
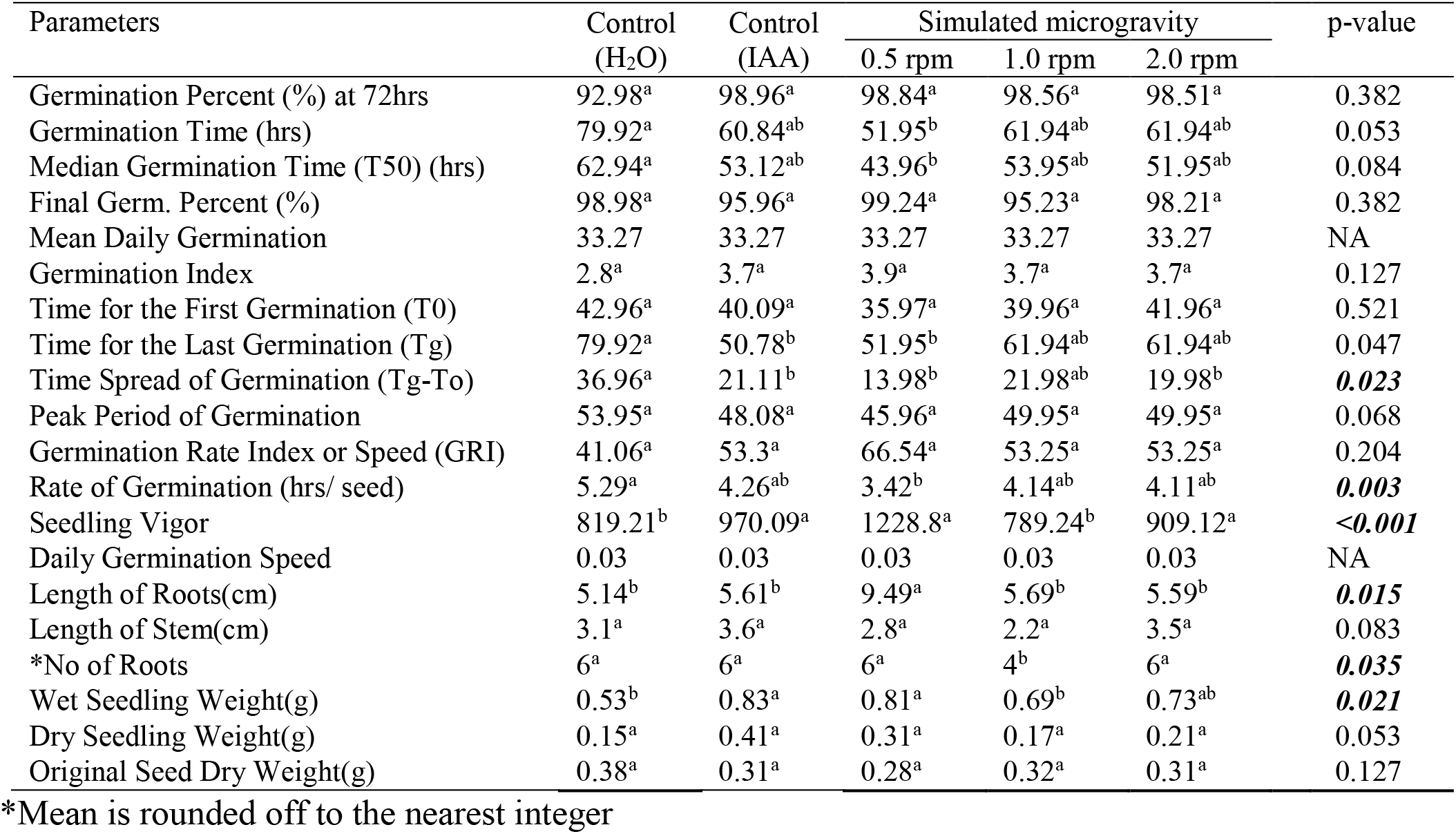
Effects indole acetic acid on the germinability and germination parameters of *Zea mays* after exposure to microgravity

| Parameters | Control<br>(H<sub>2</sub>O) | Control<br>(IAA) | Simulated microgravity |  |  | p-value |
| --- | --- | --- | --- | --- | --- | --- |
|  |  |  | 0.5 rpm | 1.0 rpm | 2.0 rpm |  |
| Germination Percent (%) at 72hrs | 92.98<sup>a</sup> | 98.96<sup>a</sup> | 98.84<sup>a</sup> | 98.56<sup>a</sup> | 98.51<sup>a</sup> | 0.382 |
| Germination Time (hrs) | 79.92<sup>a</sup> | 60.84<sup>ab</sup> | 51.95<sup>b</sup> | 61.94<sup>ab</sup> | 61.94<sup>ab</sup> | 0.053 |
| Median Germination Time (T50) (hrs) | 62.94<sup>a</sup> | 53.12<sup>ab</sup> | 43.96<sup>b</sup> | 53.95<sup>ab</sup> | 51.95<sup>ab</sup> | 0.084 |
| Final Germ. Percent (%) | 98.98<sup>a</sup> | 95.96<sup>a</sup> | 99.24<sup>a</sup> | 95.23<sup>a</sup> | 98.21<sup>a</sup> | 0.382 |
| Mean Daily Germination | 33.27 | 33.27 | 33.27 | 33.27 | 33.27 | NA |
| Germination Index | 2.8<sup>a</sup> | 3.7<sup>a</sup> | 3.9<sup>a</sup> | 3.7<sup>a</sup> | 3.7<sup>a</sup> | 0.127 |
| Time for the First Germination (T0) | 42.96<sup>a</sup> | 40.09<sup>a</sup> | 35.97<sup>a</sup> | 39.96<sup>a</sup> | 41.96<sup>a</sup> | 0.521 |
| Time for the Last Germination (Tg) | 79.92<sup>a</sup> | 50.78<sup>b</sup> | 51.95<sup>b</sup> | 61.94<sup>ab</sup> | 61.94<sup>ab</sup> | 0.047 |
| Time Spread of Germination (Tg-To) | 36.96<sup>a</sup> | 21.11<sup>b</sup> | 13.98<sup>b</sup> | 21.98<sup>ab</sup> | 19.98<sup>b</sup> | **0.023** |
| Peak Period of Germination | 53.95<sup>a</sup> | 48.08<sup>a</sup> | 45.96<sup>a</sup> | 49.95<sup>a</sup> | 49.95<sup>a</sup> | 0.068 |
| Germination Rate Index or Speed (GRI) | 41.06<sup>a</sup> | 53.3<sup>a</sup> | 66.54<sup>a</sup> | 53.25<sup>a</sup> | 53.25<sup>a</sup> | 0.204 |
| Rate of Germination (hrs/ seed) | 5.29<sup>a</sup> | 4.26<sup>ab</sup> | 3.42<sup>b</sup> | 4.14<sup>ab</sup> | 4.11<sup>ab</sup> | **0.003** |
| Seedling Vigor | 819.21<sup>b</sup> | 970.09<sup>a</sup> | 1228.8<sup>a</sup> | 789.24<sup>b</sup> | 909.12<sup>a</sup> | **<0.001** |
| Daily Germination Speed | 0.03 | 0.03 | 0.03 | 0.03 | 0.03 | NA |
| Length of Roots(cm) | 5.14<sup>b</sup> | 5.61<sup>b</sup> | 9.49<sup>a</sup> | 5.69<sup>b</sup> | 5.59<sup>b</sup> | **0.015** |
| Length of Stem(cm) | 3.1<sup>a</sup> | 3.6<sup>a</sup> | 2.8<sup>a</sup> | 2.2<sup>a</sup> | 3.5<sup>a</sup> | 0.083 |
| *No of Roots | 6<sup>a</sup> | 6<sup>a</sup> | 6<sup>a</sup> | 4<sup>b</sup> | 6<sup>a</sup> | **0.035** |
| Wet Seedling Weight(g) | 0.53<sup>b</sup> | 0.83<sup>a</sup> | 0.81<sup>a</sup> | 0.69<sup>b</sup> | 0.73<sup>ab</sup> | **0.021** |
| Dry Seedling Weight(g) | 0.15<sup>a</sup> | 0.41<sup>a</sup> | 0.31<sup>a</sup> | 0.17<sup>a</sup> | 0.21<sup>a</sup> | 0.053 |
| Original Seed Dry Weight(g) | 0.38<sup>a</sup> | 0.31<sup>a</sup> | 0.28<sup>a</sup> | 0.32<sup>a</sup> | 0.31<sup>a</sup> | 0.127 |

\*Mean is rounded off to the nearest integer

As previously mentioned, and similarly to gibberellic acid and indole acetic acid, the effect of ascorbic acid on germination percentage in microgravity-exposed seeds has been presented (Table3). The final germination percentage ranged from 94.16 to 98.98 percent, with a p>0.05. While the results in Figure 1 showed that exposing seeds to microgravity for up to 2.0 rpm significantly delayed germination, the results in table 3 showed that exposing seeds that had been first soaked in ascorbic acid to microgravity for up to 2.0 rpm did not significantly delay germination when compared to the control; in this case, the time taken for first germination was reported to be between 31.97-42.98hrs (p>0.05). The time spread of germination, or the time during which seeds germinated, was 36.96 hrs in the control, 18.77 hrs when seeds were first exposed to vitamin C but not to microgravity, and 11.99-19.98 hrs when seeds were first soaked in ascorbic acid and then exposed to simulated microgravity. There were no major variations in seedling dry weight regardless of the degree of simulated microgravity, as seedling dry weight ranged from 0.11g in 2.0 rpm to 0.25g in 1.0 rpm (p>0.05), as reported in Tables 1 and 2.

**Plate 2:**
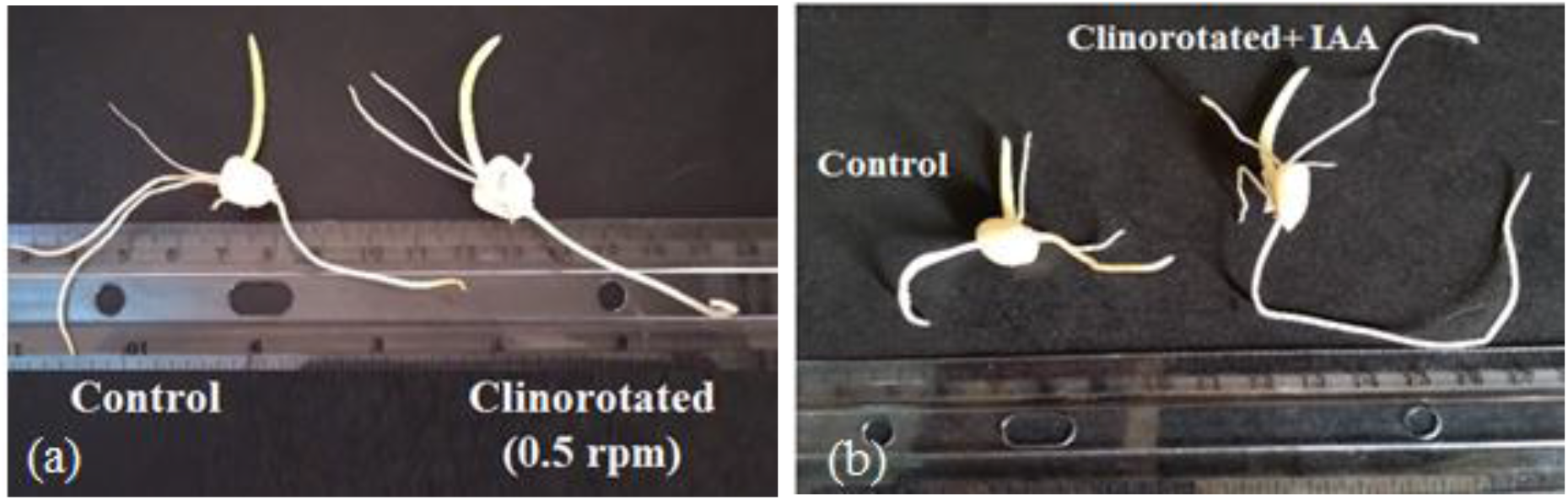
(a) Clinorotated seedling and (b) IAA-primed clinorotated maize seedling compared with the control at 50 hrs after germination initiation

**Table 3:** Effects ascorbic acid on the germinability and germination parameters of *Zea mays* after exposure to microgravity

| Parameters | Control<br>(H<sub>2</sub>O) | Control<br>(Vit. C) | Simulated microgravity |  |  | p-value |
| --- | --- | --- | --- | --- | --- | --- |
|  |  |  | 0.5 rpm | 1.0 rpm | 2.0 rpm |  |
| Germination Percent (%) at 72hrs | 98.98<sup>a</sup> | 94.16<sup>a</sup> | 96.87<sup>a</sup> | 98.32<sup>a</sup> | 97.46<sup>a</sup> | 0.468 |
| Germination Time (hrs) | 79.92<sup>a</sup> | 63.42<sup>ab</sup> | 49.95<sup>b</sup> | 51.95<sup>b</sup> | 54.95<sup>b</sup> | **<0.001** |
| Median Germination Time (T50) (hrs) | 62.94<sup>a</sup> | 51.25<sup>ab</sup> | 43.96<sup>b</sup> | 39.96<sup>b</sup> | 41.96<sup>b</sup> | **0.005** |
| Final Germ. Percent (%) | 98.98<sup>a</sup> | 94.16<sup>a</sup> | 96.87<sup>a</sup> | 98.32<sup>a</sup> | 97.46<sup>a</sup> | 0.468 |
| Mean Daily Germination | 33.27 | 33.27 | 33.27 | 33.27 | 33.27 | NA |
| Germination Index | 2.8<sup>a</sup> | 3.7<sup>a</sup> | 3.8<sup>a</sup> | 3.9<sup>a</sup> | 3.9<sup>a</sup> | 0.137 |
| Time for the First Germination (T0) | 42.96<sup>a</sup> | 42.98<sup>a</sup> | 37.96<sup>a</sup> | 31.97<sup>a</sup> | 35.97<sup>a</sup> | 0.329 |
| Time for the last Germination (Tg) | 79.92<sup>a</sup> | 63.22<sup>ab</sup> | 49.95<sup>b</sup> | 51.95<sup>b</sup> | 54.95<sup>b</sup> | **<0.001** |
| Time Spread of Germination (Tg-To) | 36.96<sup>a</sup> | 18.77<sup>ab</sup> | 11.99<sup>b</sup> | 19.98<sup>ab</sup> | 18.98<sup>b</sup> | 0.058 |
| Peak Period of Germination | 53.95<sup>a</sup> | 50.92<sup>a</sup> | 37.96<sup>b</sup> | 34.97<sup>b</sup> | 37.96<sup>b</sup> | 0.038 |
| Germination Rate Index or Speed (GRI) | 41.06<sup>b</sup> | 43.3<sup>b</sup> | 63.24<sup>ab</sup> | 73.23<sup>a</sup> | 73.23<sup>a</sup> | **0.004** |
| Rate of Germination (hrs/ seed) | 5.29<sup>a</sup> | 4.01<sup>ab</sup> | 3.33<sup>b</sup> | 3.52<sup>b</sup> | 3.78<sup>b</sup> | **<0.001** |
| Seedling Vigor | 819.21<sup>a</sup> | 790.02<sup>ab</sup> | 999.03<sup>a</sup> | 539.48<sup>b</sup> | 999.03<sup>a</sup> | **0.027** |
| Daily Germination Speed | 0.03 | 0.03 | 0.03 | 0.03 | 0.03 | NA |
| Length of Roots(cm) | 5.12<sup>b</sup> | 4.81<sup>bc</sup> | 8.99<sup>a</sup> | 3.53<sup>c</sup> | 7.19<sup>a</sup> | **<0.001** |
| Length of Stem(cm) | 3.1<sup>a</sup> | 3.1<sup>a</sup> | 1.1<sup>b</sup> | 1.9<sup>b</sup> | 2.8<sup>ab</sup> | **0.033** |
| *No of Roots | 6<sup>ab</sup> | 5<sup>b</sup> | 5<sup>b</sup> | 5<sup>b</sup> | 8<sup>a</sup> | **0.014** |
| Wet Seedling Weight(g) | 0.53<sup>a</sup> | 0.64<sup>a</sup> | 0.71<sup>a</sup> | 0.78<sup>a</sup> | 0.76<sup>a</sup> | 0.283 |
| Dry Seedling Weight(g) | 0.15<sup>a</sup> | 0.12<sup>a</sup> | 0.22<sup>a</sup> | 0.25<sup>a</sup> | 0.11<sup>a</sup> | 0.138 |
| Original Seed Dry Weight(g) | 0.38<sup>a</sup> | 0.29<sup>a</sup> | 0.32<sup>a</sup> | 0.29<sup>a</sup> | 0.32<sup>a</sup> | 0.562 |

\*Mean is rounded off to the nearest integer

The asymptotic significance values resulting from the comparison of the impact of these growth stimulators on the germination of maize have been provided in Table 4 in an attempt to compare the capacity of the three growth stimulators to ameliorate the influence of microgravity on seed germination. There were no major variations in the ability of the three growth stimulators to influence germination percentage or time taken for first and last germination under the microgravitational impact attributed to clinorotation at 0.5 rpm, according to the results. However, indole acetic acid improved seedling vigor better than gibberellic acid and ascorbic acid under microgravity influence at 0.5 rpm. Similarly, under microgravitational impact at 0.5 rpm, indole acetic acid had a stronger influence on root length in maize germinants. Gibberellic acid, rather than indole acetic acid or ascorbic acid, most likely influences germination time at 1.0 rpm. Gibberellic acid is also more likely than the other two growth stimulants to affect the peak time and rate of germination. Table 4 also showed that indole acetic acid and ascorbic acid improved seedling vigor more than gibberellic acid when microgravity was reached at 2.0 rpm in the clinostat.

**Table 4:**
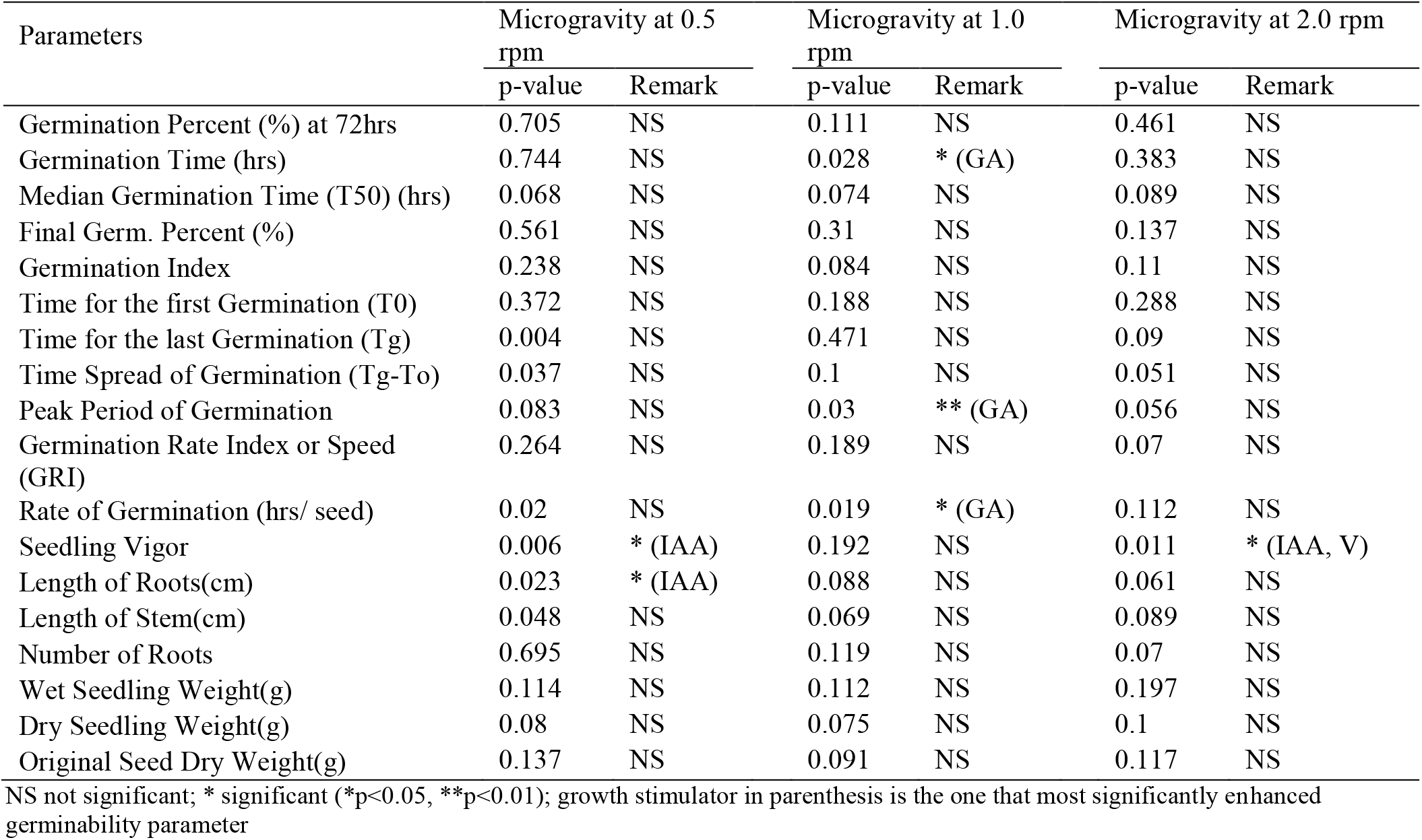
Asymptotic significance values arising from comparison of the effects of gibberellic (GA), indole acetic (IAA) and ascorbic acids on the germination parameters of *Zea mays* influenced by each simulated microgravity

In comparison to control seeds sown under normal gravity, Table 5 indicates percentage differences in time to first germination and final seedling dry weight at 72 hrs. The results show that under simulated gravity, there was a substantial reduction in time to first germination by as much as 44.9 percent due to the influence of gibberellic acid, indole acetic acid, and ascorbic acid, implying that the growth stimulators’ influence accelerated germination time. When seedling dry weight was also examined, the results showed that when microgravity-exposed seeds were handled with gibberellic acid, indole acetic acid, and ascorbic acid for 72 hrs, seedling dry weight increased significantly (Table 5). Seedling dry weight increased by 198.1 percent under the influence of gibberellic acid at 1.0 rpm, and by 201.2 percent under the influence of indole acetic acid at 2.0 rpm (Table 5). The implication is that exposing germinating seeds to microgravity would boost their growth statistics if they were earlier primed with gibberellic acid, indole acetic acid, and ascorbic acid beforehand.

**Table 5:** Percentage changes in time to first germination and final seedling dry weight at 72 hrs compared to control seeds under gravity

| Plant growth stimulator | Simulated microgravity |  |  |
| --- | --- | --- | --- |
|  | 0.5 rpm | 1.0 rpm | 2.0 rpm |
| ***Δ% Time to first germination*** |  |  |  |
| 150 ppm GA | -1.3 | -25.4 | -37.1 |
| 150 ppm IAA | -23.2 | -30.7 | -26.3 |
| 150 ppm Vit. C | -16.7 | -44.9 | -37.1 |
| ***Δ% Seedling dry wt.*** |  |  |  |
| 150 ppm GA | 128.1 | 198.1 | 165.1 |
| 150 ppm IAA | 31.6 | 63.5 | 201.2 |
| 150 ppm Vit. C | -12.3 | 101.9 | 32.5 |

***Δ% percentage change***
*Values presented are percentage changes in time to first germination and final seedling dry weight at 72 hrs compared to control seeds under gravity. Whereas negative values indicate percentage reductions while positive values indicate positive increase in the measured parameter compared to control seeds under gravity.*

## DISCUSSION

Before maize seed stimulation with plant growth promoters, the reduced germination parameters observed in the current study when seeds were exposed to microgravity as against the control indicated the perturbations of many biological phenomena due to the effect of altered gravity (De Micco et al., 2013). This is consistent with the work of Orukpe et al. (2021) where he explained that microgravity is a major impediment to maize seed germination in most parts of the world. In order to ameliorate this effect, the researchers discovered that exposing seeds to growth stimulants or seed pre-treatment improved seed germination rate, seedling vigor, and growth statistics, reducing the effect of microgravity on maize seed germinability (p>0.05). Despite the fact that responses differed with different growth stimulators, the three growth stimulators used (Indole acetic acid, Gibberellic acid, and Ascorbic acid) had different effects on maize seed germinability. GA improved germination by extending the peak time and speed of germination, while IAA improved root length and seedling vigor. Ascorbic acid’s most notable result was an improvement in seedling vigor. In the clinostat, indole acetic acid and ascorbic acid increased seedling vigor more than gibberellic acid since microgravity was achieved at 2.0 rpm.

The three distinct phases of seed germination are imbibition, lag phase, and radicle growth and emergence (Bradford, 1995). Zabel et al. (2016) explained that germination is needed for plant growth and development. If a plant survives, that is because it was the first to germinate. Seed pre-treatment significantly increased final germination percent, median germination period, peak germination length, germination time, and seedling vigor of maize, even under the influence of simulated microgravity, according to the findings of this study and also consistent with the work of Musa and Ikhajiagbe (2021) and Mshelbula *et al*. (2015). Priming seeds with growth hormones boosts seed and seedling vigor through metabolic and biochemical processes that occur during regulated hydration followed by dehydration (Becerra-Vázquez et al., 2020).

The current study showed that there is no significant effects of the growth regulators used in the current study on the number of roots. This is consistent with the work of Hoson et al. (1992) when they looked at how clinorotation affected cress (*Lepidium sativum*), pea (*Pisum sativum*) and azuki bean (*Vigna angularis*) which showed no significant difference on roots number. The rise in seed germination as a result of seed priming is consistent with Murungu et al. (2004) and Basra et al (2006). GA had a greater effect on germination rate at all concentrations as compared to other priming reagents. This is likely because GA have been implicated in the regulation of many plant growth and developmental processes, with an emphasis on stem elongation (Hedden and Phillips 2000; Sevik and Guney (2013); Zhang et al (2007). The rapid utilization of GA-treated seeds in the synthesis of various amino acids and amides (Gupta and Mukherjee, 1982) is the explanation for the increased germination time.

The higher seed dry weight observed in the control as against the microgravity may be because the microgravity perturbation disrupt the water use efficiency of the seeds even under growth stimulators. Because the clinostat movement improves seed drying which affects seed weight (Bradford, 2006). Since the microgravity condition is also a stress factor for plants, according to this report. Microgravity has the potential to disrupt plant growth processes, but with the use of growth stimulators, the effect of the stress exerted by microgravity is minimized (Cowles et al., 2008) encouraging the possibilities of growing maize seeds even during space exploration (Ikhajiagbe et al., 2007)

## CONCLUSION

Since the findings of this research have shown the possibilities of using plant growth stimulators such as IAA, GA and AA to improve maize germination parameters even under microgravity condition, this showed the novelty of this research. With this, idea as to what extent we can prepare for future space exploration can be predicted. Further research is encouraged to investigate the influence of other plant growth stimulators and also to test it on other important cereals such as rice.

## Acknowledgment

Special appreciation to the Space-Earth Environment Research Laboratory, University of Benin, Edo State, Nigeria, for permission to use their facility during the study. This study was privately funded.

